# Network Insights into Improving Drug Target Inference Algorithms

**DOI:** 10.1101/2020.01.17.910885

**Authors:** Muying Wang, Heeju Noh, Ericka Mochan, Jason E. Shoemaker

**Affiliations:** Department of Chemical and Petroleum Engineering, Swanson School of Engineering, University of Pittsburgh, Pittsburgh, Pennsylvania, USA; Department of Systems Biology, Columbia University, New York, NY 10032, USA; Institute for Chemical and Bioengineering, ETH Zurich, Zurich 8093, Switzerland; Department of Mathematics and Data Analytics, Carlow University, Pittsburgh, PA 15213, USA; Department of Computational and Systems Biology, School of Medicine, University of Pittsburgh, Pittsburgh, PA 15213, USA; The McGowan Institute for Regenerative Medicine, Pittsburgh, PA 15213, USA

**Keywords:** drug target inference, network topology, gene expression, protein-protein interactions, betweenness centrality

## Abstract

To improve the efficacy of drug research and development (R&D), a better understanding of drug mechanisms of action (MoA) is needed to improve drug discovery. Computational algorithms, such as ProTINA, that integrate protein-protein interactions (PPIs), protein-gene interactions (PGIs) and gene expression data have shown promising performance on drug target inference. In this work, we evaluated how network and gene expression data affect ProTINA’s accuracy. Network data predominantly determines the accuracy of ProTINA instead of gene expression, while the size of an interaction network or selecting cell/tissue-specific networks have limited effects on the accuracy. However, we found that protein network betweenness values showed high accuracy in predicting drug targets. Therefore, we suggested a new algorithm, TREAP (https://github.com/ImmuSystems-Lab/TREAP), that combines betweenness values and adjusted *p*-values for target inference. This algorithm has resulted in higher accuracy than ProTINA using the same datasets.

## Introduction

The innovation of treatments for diseases remains a challenging task [1-6]. The efficiency of pharmaceutical research and development (R&D), quantified by the number of new drugs per billion US dollars spent, dramatically declined from 1950 to 2010 [2]. A large group of drug candidates fail in clinical trials because they are not effective or safe in humans [2, 5, 7-9]. A major reason is that the systematic effects of drug candidates are not well studied or modelled in the drug discovery process, and a better understanding of their mechanisms of action (MoA) can help improve the efficiency of drug R&D [2, 10-12].

Two types of computational approaches have been reported to study MoAs of drugs by modeling high-throughput biological data: comparative analysis and network-based algorithms [13, 14]. Comparative analysis approaches, such as the Connectivity Map [15], have been used to predict molecular targets of drugs and assist in drug repurposing [15-20]. They utilize expression profiles as drug signatures and compare with drugs having known targets, assuming that drugs with high similarities share the same targets. These approaches much rely on prior knowledge of drugs, thus have limitations in predicting de novo targets.

Network-based algorithms predict drug or disease targets by combining network information and transcriptomic data [14, 21-27]. Two recent representatives, DeMAND [22] and ProTINA [14], model the systemic dysregulation of regulatory network caused by a drug treatment, connecting molecular interactions with differential expression (DE). The regulatory network is generated by using protein-protein interactions (PPIs) and protein-gene interactions (PGIs) obtained from self-curated or public databases, such as STRING [28] and CellNet [29]. Similar to what has been reported by Noh et al [14], our preliminary research showed that ProTINA outperforms DeMAND when tested by the same gene expression and network datasets (Figure S1). Therefore, in this work, we focused on studying ProTINA’s performance.

For ProTINA, a regulatory network, directing from proteins or transcriptional factors (TFs) to regulated genes, is generated from input PPIs and PGIs based on certain rules (Figure 1) [14]. The assumption is that the log fold change (LFC) of a gene is the linear combination of the LFCs of all proteins and TFs that regulate it. The weights are computed by linear regression methods and then integrated into a score for each regulator, a protein or a TF. Different from DeMAND, ProTINA may result in negative or positive scores, representing attenuation or enhancement, respectively. Regulators of larger magnitudes are more likely to be targets (red nodes in Figure 1). Showing promising results in predicting *in vitro* datasets, DeMAND and ProTINA have provided a new direction in identifying drug targets and toxicity [14, 22].

**Figure 1.**
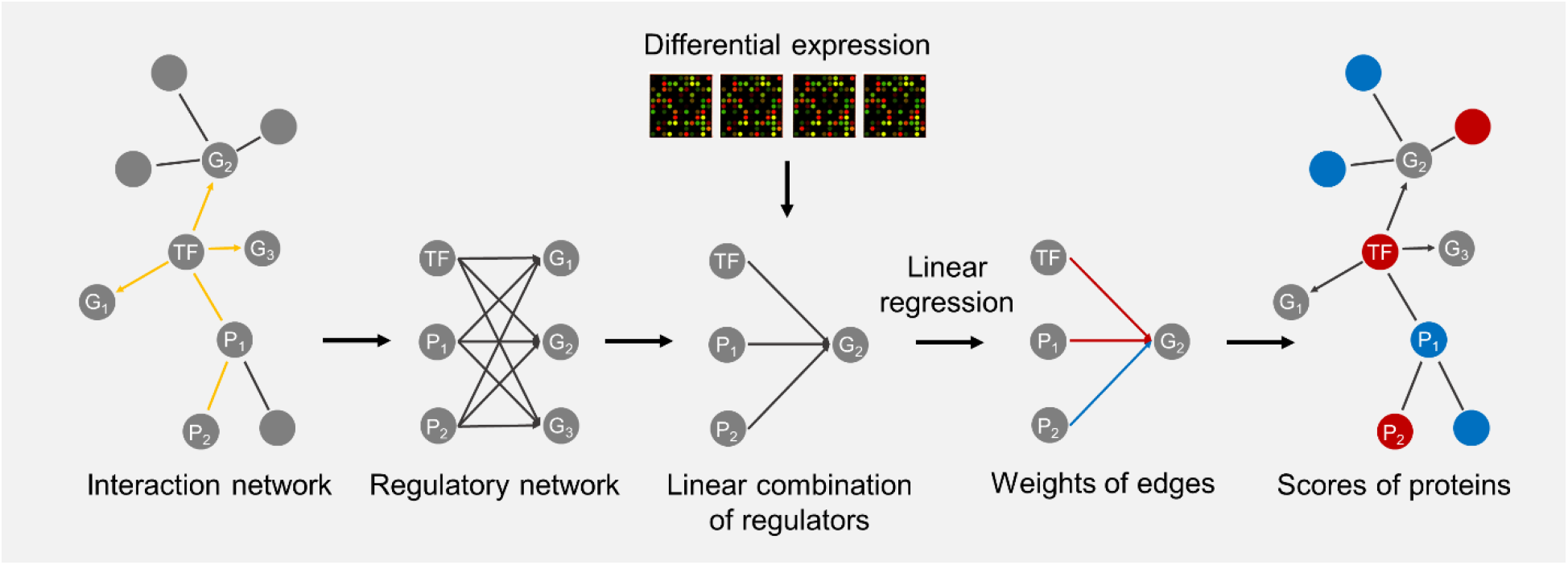
An overview of ProTINA algorithm. Each node refers to a transcription factor (TF), a non-TF protein (P) or a gene (G). Arrows present the directions of interactions or edges. The significance of an edge or protein (including TFs) is color coded, where red refers to high significance while blue refers to low significance.

However, as target inference algorithms become more complicated, it is unclear what roles gene expression and network data play. A recent study has shown that an accurate description of network topology is able to cover 65% of the perturbation patterns predicted by a full biochemical model with kinetic parameters [30]. Several studies have shown that proteins associated with disease and proteins that are drug targets have significantly different positions within biological networks [31-34]. For target inference algorithms, it remains an open question as to which kind of biological data most affects the accuracy. Furthermore, algorithms can infer drug targets in a cell/tissue type-specific manner [14, 22, 27], and it is unknown how efficient or meaningful cell/tissue type-specific network data is for target inference. Answering these questions can provide us with insights into future algorithm improvement.

In this work, we evaluated the impact of gene expression and network data, using the human B cell microarray data from the DREAM challenge (referred to as DP14) as our benchmark dataset [35], and introduced a new algorithm to predict drug targets. Firstly, we found that ProTINA’s scores are mostly determined by network data through permutation tests on gene expression. Secondly, we tested how the selection of networks affects prediction accuracy. Surprisingly, the effects of size or cell type are negligible.Next, our analysis suggested that network betweenness values can accurately predict drug targets. The performance is comparable with ProTINA and is consistent regardless of the network size. Lastly, we proposed TREAP to combine betweenness values and adjusted *p*-values from DE for target inference, which has outperformed ProTINA in accuracy. Moreover, the simplicity of the algorithm makes it more tractable to users who are not experts in systems and network biology. Our future work will focus on better balancing both types of data and trying other methods, such as machine learning, to improve prediction accuracy.

## Materials and Methods

### Gene expression data used in the analysis

The microarray data of human Diffuse Large B-Cell Lymphoma (DLBCL) OCI-LY3 cell line treated with 14 different drugs under diverse doses at 3 time points, 6, 12 and 24 hours post treatment were obtained from the NCI-DREAM challenge drug synergy dataset, DP14 (GEO accession: GSE51068) [35]. Three samples treated with ‘Aclacinomycin A’ under a lower dose were dropped due to less significance. The microarray data of human liver cell line HepG2 treated with 62 genotoxic or non-genotoxic chemicals at 12, 24 and 48 hours post treatment were obtained from literature, referred to as HepG2 in this work (GEO accession: GSE28878) [36]. The microarray data of mouse pancreatic cells treated with 29 chromatin-targeting compounds were also obtained from GEO database, referred to as MP (GEO accession: GSE36379) [37]. For all three datasets, raw data were normalized using the RMA function from the “*affy*” R package [38]. The log2 fold change (LFC) values and Benjamini–Hochberg adjusted *p*-values (adjusted *p*-values) were calculated by the “*limma*” R package [39]. Probes were mapped to gene symbols by using the “*hgu219.db*” R package for human microarray data and “*moe430a.db*” for mouse data. Those with the lowest average BH-adjusted *p*-value across all samples were chosen when multiple probes were mapped to the same gene.

### Networks used in the analysis and calculation of topological features

Human or mouse PPIs and their associated confidence scores were obtained from the STRING database [28]. Interactions with experimental proof or from curated databases (the channels of ‘experiments’ and ‘databases’) were extracted. Interactions transferred from other species or duplicated entries were excluded. Subnetworks were obtained by applying thresholds ranging from 0.4 to 0.9 to the PPI network, referred to as PPI04, PPI05, PPI06, PPI07, PPI08 and PPI09, respectively in this paper.

Human PGIs and their confidence scores were obtained from the Regulatory Circuits, a database of predicted, cell/tissue type-specific PGIs [40]. PGI networks of 8 different cell/tissue types were studied in this analysis: ‘lymphocytes of B lineage’, ‘lymphocytes’, ‘lymphoma’, ‘myeloid leukocytes’, ‘lung’, ‘heart’, ‘epithelial cells’ and ‘hepatocellular carcinoma cell line’. The network of ‘lymphocytes of B lineage’ were predicted by samples including those from DLBCL, the same cell line with DP14 [40], thus was chosen as a reference for analysis of DP14. PGI subnetworks for each cell/tissue type, namely PGI05, PGI10, PGI15 and PGI20, were obtained by thresholds ranging from 0.05 to 0.20, respectively. Mouse PGIs were compiled from two manually curated databases of transcriptional regulatory networks: TRRUST (version 2) [41] and RegNetwork [42]. These interactions are not cell/tissue type-specific, and no threshold was applied to them prior to analysis of MP.

Degree or betweenness values were calculated by the “*igraph*” R package [43]. PPIs or the combination of PPIs and PGIs (PPI+PGI) were treated as undirected graphs, while PGIs were treated as directed graphs.

### Reference drug targets

The reference targets of each chemical were extracted from STITCH database (version 5.0) [44, 45] for analyses of DP14, HepG2 and MP. Only targets with experimental proof or from curated databases were collected as shown in Table S1.

### Prediction of drug targets by ProTINA

LFC values, PPI and PGI subnetworks were analyzed by “*protina*” R package [14]. Slope matrices of each time point were calculated following the user manual. For samples with only two timepoints, control samples served as 0hr post treatment to calculate associated slope matrices. Samples from different doses for the same drug were treated as separate groups.

### Prediction of drug targets by TREAP

For target prediction by TREAP, the assumption is that genes with high betweenness values or low adjusted *p*-values are more likely to be drug targets. Adjusted *p*-values and PPI+PGI betweenness values were calculated as explained in the former sections. Ranks of genes were obtained by sorting betweenness values and adjusted *p*-values, respectively, and genes with the same betweenness or adjusted *p*-value shared the same rank. Final scores were calculated by summing up the ranks from both metrics for each gene. In this work, all analyses on TREAP used 0.9 as the threshold for human or mouse PPIs and 0.20 for human PGIs. No threshold was applied to mouse PGIs.

### Calculation and comparison of AUROC values

Area under the receiving operator characteristics (AUROC) values in this paper were calculated by comparing scored proteins with reference drug targets through the “*pROC*” R package [46]. As ProTINA scores can be positive or negative, the absolute scores were used to calculate AUROC. The median AUROC across all drugs in each dataset was calculated to represent accuracy of a whole test. For drugs having more than one doses, the AUROC values of low doses were excluded. In terms of topological features, degree or betweenness values were directly used to calculate AUROC values without pre-processing. TREAP scores were directly used for calculation of AUROC without preprocessing. Difference in any pair of chosen tests were computed by performing pairwise *t*-test between their AUROC values. A *p*-value less than 0.05 were regarded as significantly different.

### Permutation tests on gene expression

The null hypothesis for permutation tests in this work is that the median AUROCs of randomized gene expression are smaller than that of nonrandomized gene expression, and the *p*-values were calculated accordingly. For ProTINA, gene labels for DP14 that refer to the rows of LFC and associated slope matrices were randomly shuffled for 1000 times. Randomized data were applied to ProTINA under the same network setup, PPI09 and PGI20. For TREAP, gene labels of adjusted *p*-values for each dataset, namely DP14, HepG2 and MP were randomly shuffled for 1000 times, respectively, and drug targets were predicted using PPI09 and PGI20 (for MP no threshold was applied). AUROC and median values were calculated as explained in the former section.

## Results

### Permutation tests show that ProTINA is predominantly determined by network data

To understand how much network or gene expression data contribute to ProTINA’s accuracy, we performed 1000 permutation tests by randomizing the LFC gene expression values (Materials and Methods). To shorten the computation time on ProTINA, the smallest PPI and PGI subnetworks (PPI09 and PGI20) were chosen for this analysis (discussed more in the following section). For prediction scores obtained from ProTINA, the area under the receiver operating characteristics (AUROC) values were calculated per drug, and the median AUROC across all drugs is used as a metric for each test.

The median AUROC obtained by nonrandomized LFCs is 0.799. As shown in Fig. 2, 220 of 1000 permutation tests have higher median AUROC values than that (one-tailed *p*-value = 0.221). Most tests have similar accuracy to the original test. This indicates that randomizing LFCs does not diminish ProTINA’s accuracy significantly, therefore, network data determines most of ProTINA’s performance.

**Figure 2.**
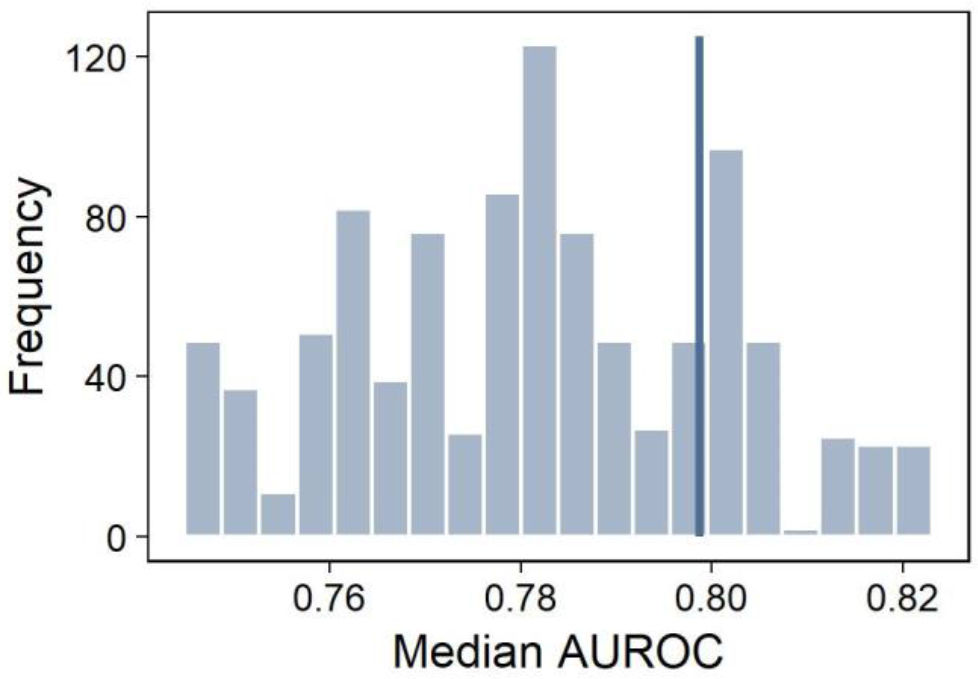
1000 Permutation tests were performed by randomizing the gene expression and calculating the median AUROCs. The blue vertical line refers to the median AUROC obtained by nonrandomized gene expression.

### Selection of networks has limited effects in the prediction accuracy of ProTINA

Next, we studied how the selection of PPIs and PGIs affects target inference accuracy. PPI and PGI subnetworks of different sizes or cell/tissue types were tested using the same gene expression data from DP14 (Materials and Methods). Similar to permutation tests, the median AUROC represents the accuracy for each PPI-PGI combination.

In total, 24 PPI-PGI combinations of different sizes were tested on ProTINA. As shown in Figure 3a, ProTINA favors small PPI and large PGI subnetworks. The combination of PPI09 and PGI05 shows the highest accuracy, and its median AUROC is 0.821. As the threshold increases from 0.4 to 0.9, the number of interactions in PPI subnetwork ranges from 380375 to 281357 (Figure S2), and the median AUROC increases for most tests. For example, the median AUROC increases from 0.785 (PPI04) to 0.811 (PPI09) for analyses under PGI10. But there are PPI subnetworks that do not follow this trend. Under PGI05, PPI05 shows lower median AUROC than PPI04 (0.784 and 0.796, respectively). For PGI subnetworks, as the number of interactions range from 123394 to 5932 (Figure S2), the median AUROC shows an opposite trend to that of PPI subnetworks (Figure 3a). An example is that the median AUROC decreases from 0.821 (PGI05) to 0.785 (PGI20) when using PPI09. Most tests show a consistent trend except for those using PPI05. A possible reason is that new proteins and associated interactions are included in the network as the threshold for PGIs changes from 0.05 to 0.10. So that the network topology is changed, and predictions from ProTINA are affected accordingly.

**Figure 3.**
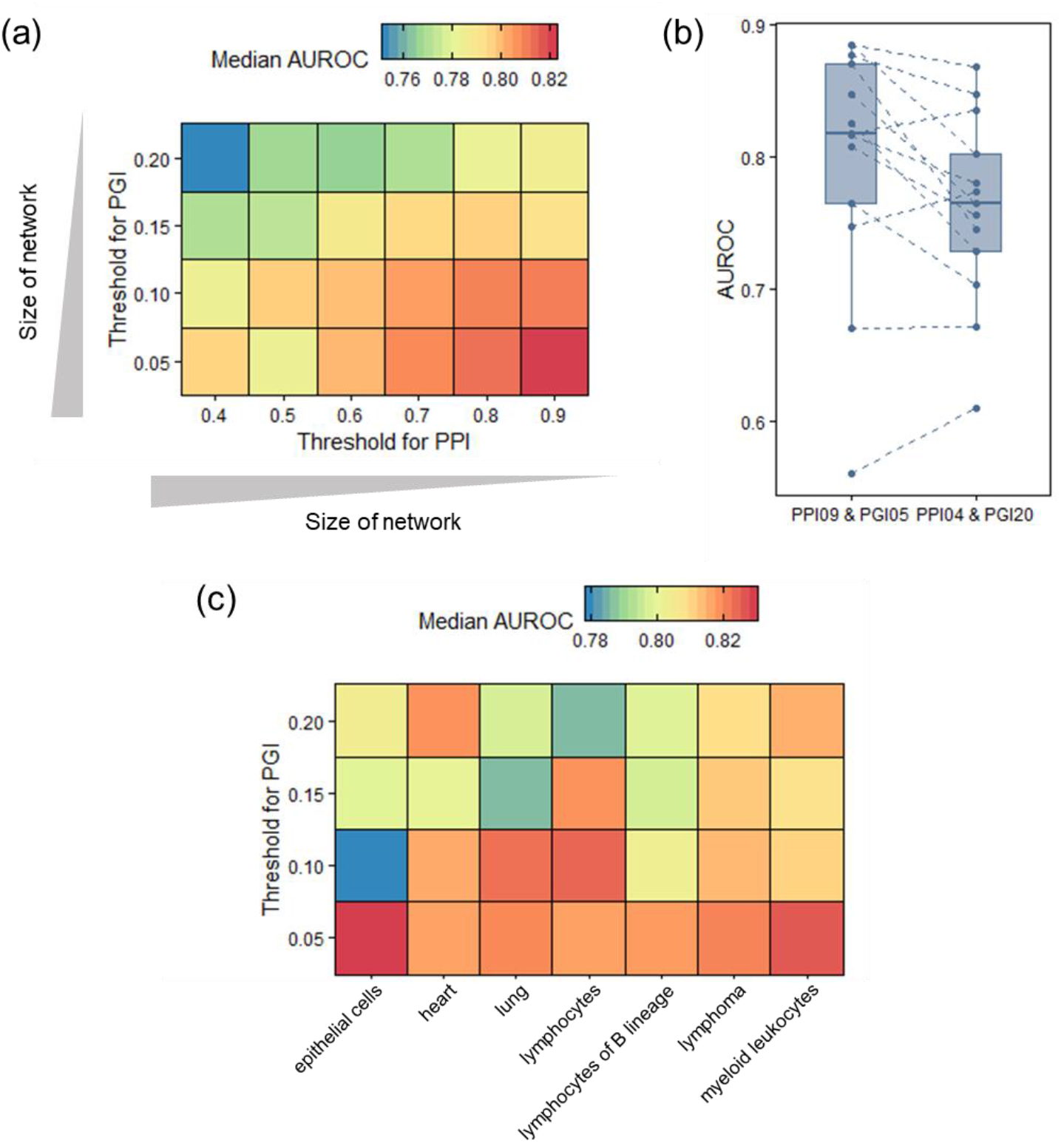
Prediction accuracy of ProTINA using networks of different sizes or cell/tissue types. (a) PPI or PGI subnetworks of different sizes were tested on ProTINA to predict targets for DP14. The axes refer to the confidence thresholds for PPI (x axis) and PGI (y axis) subnetworks, and the median AUROC values are the metric for prediction accuracy. Among all PPI-PGI subnetwork combinations, PPI09-PPI05 and PPI04-PPI20 have the highest and lowest accuracy in terms of median AUROC values, respectively. Panel (b) shows the boxplot of these two groups. Each dot represents the AUROC of a drug. (c) PGI subnetworks of 7 cell/tissue types were applied to ProTINA for target prediction.

However, median AUROC values vary in a small range while the size of PPI or PGI subnetworks changes significantly (Figure 3a, 3b and Figure S3). The highest and lowest median AUROC values for ProTINA are 0.821 and 0.753, with a difference less than 0.1 (Figure 3b). In addition, most of these differences are insignificant. When comparing with the test using PPI09 and PGI20, none of the 24 tests are significant in AUROC values (*p*-value > 0.05 for all tests), although the combination of PPI04 and PGI15 has resulted in a pairwise *p*-value of 0.057. Furthermore, we studied the effects on standard deviations (SDs) of AUROC values across 12 drugs for each test. All of them maintain at a low level below 0.13. In summary, we conclude that the size of networks has limited effects on the prediction accuracy of ProTINA, while small PPI and large PGI networks tend to improve the accuracy.

To analyze the performance of cell/tissue type-specific networks, 28 tests using PGIs from 7 cell/tissue types were performed on ProTINA using the same PPI subnetwork, PPI09. Most tests counterintuitively show similar median AUROC values regardless of cell/tissue types (Figure 3c, Figure S4). In theory, the PGI subnetworks for immune cells should have higher accuracy than non-immune cell types, and those for ‘lymphocytes of B cell lineage’ should outperform other immune cells. This is because that samples from DLBCL, the same cell line with DP14, were used to predict the interactions for ‘lymphocytes of B lineage’ [40] (Materials and Methods). However, using the AUROC values from PGI20 for ‘lymphocytes of B cell lineage’ as a reference, no other cell/tissue types are significantly different (pairwise *p*-values > 0.05 for all) under the same network setup. In conclusion, we have shown that the selection of PPIs or PGIs in terms of either the size or cell/tissue type is not the key factor to prediction accuracy of ProTINA.

### Topological features have similar prediction accuracy to ProTINA, and protein betweenness outperforms degree

Our findings have shown that ProTINA depends on network topology more than gene expression, and that it has consistent performance regardless of the network size or the cell/tissue type the network represents. These suggest that ProTINA is probably determined by some network topological feature that remains relatively stable across different PPI or PGI subnetworks, such as protein degree or betweenness. The degree of a protein is the number of proteins/genes with which it interacts, while the betweenness is a measure of bottleneckedness, e.g. the amount of information flowing through the proteins that connect the rest of the network. Analyses of these features and their effects on drug target prediction may provide meaningful insights on improving prediction accuracy.

To test our hypotheses, we studied degree and betweenness values for PPIs, PGIs and PPI-PGI combinations (referred to as PPI+PGI in the following text). Firstly, for PPIs, we compared scores obtained from ProTINA (using PPI09 and PGI20) with their associated protein degrees or betweenness values in PPI09 for each drug. The majority of the drugs show a weak but evident correlation between absolute ProTINA scores and protein degrees, however, the correlation for betweenness is much lower (Table S2). For instance, the correlation coefficient is 0.211 for ‘Rapamycin’ (Figure 4a), while the correlation of betweenness values is smaller than that of degrees, which is 0.085 for the same drug (Figure 4b). Notice that a large portion of the top 100 proteins scored by ProTINA (red points in Figure 4a, b) lie in the group of high degree or betweenness values.

**Figure 4.**
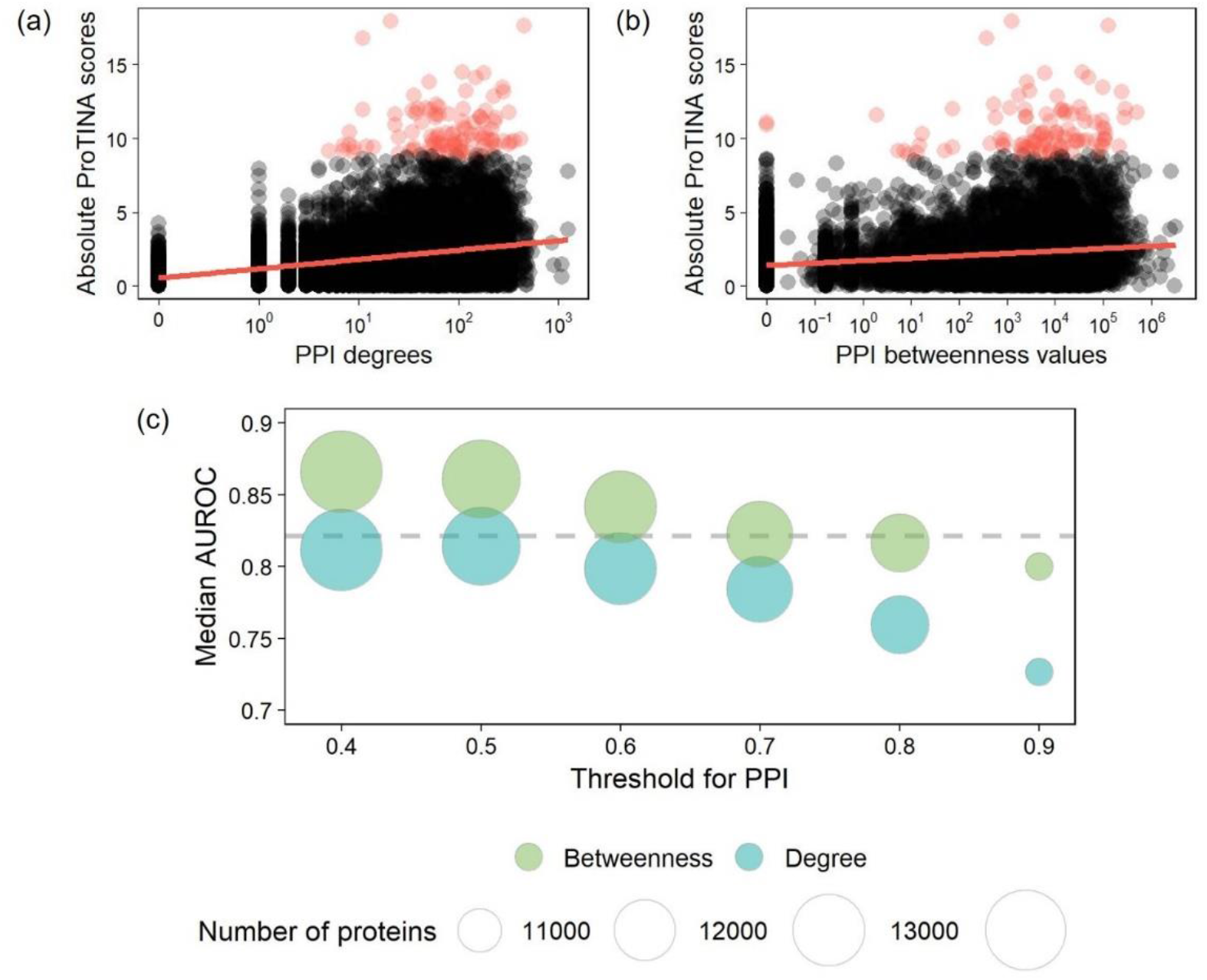
(a) Degree or (b) betweenness values of proteins in PPIs were compared with associated ProTINA scores for ‘Rapamycin’. The correlation coefficient is 0.211 for absolute ProTINA scores and degrees, and that for betweenness values is 0.085. Red points refer to the top 100 proteins scored by ProTINA. (c) The degree and betweenness values were used to predict drug targets assuming higher scores are more likely to be targets. Each point shows the median AUROC value and the number of proteins under a PPI threshold. For reference, the grey dashed line refers to the highest median AUROC achieved by ProTINA, which was obtained from using PPI09 and PGI05.

We next tried to predict drug targets by using PPI degree or betweenness values without considering gene expression or PGIs. The assumption is that proteins with higher degree or betweenness values are more likely to be targets. As shown in Figure 4c, the median AUROC values for PPI degree or betweenness values are close to those for ProTINA. What’s more, betweenness values perform better than degrees. The highest and lowest median AUROC values for degrees are 0.814 and 0.727 (Figure 4c), and those for betweenness values are 0.866 and 0.800, even higher than associated median AUROC values for ProTINA under the same network setup. As the size of PPI subnetworks shrinks, the median AUROCs for degrees decreases accordingly (Figure 4c), with a correlation coefficient value of -0.950 between the medians and thresholds. While the decrease of network size also diminishes the accuracy of betweenness, the median AUROCs remain higher than those of degrees and decrease relatively slower.

Secondly, the degree and betweenness values of PGI subnetworks were also compared with associated ProTINA scores, however, there are no clear trends between them (Table S2). In addition, drug target prediction based on PGI degree or betweenness values are not comparable with that by PPI topological features (Figure S5). This might be related to the limited amount of PGI interactions.

Lastly, we calculated topological features for PPI+PGI and compared them with ProTINA scores. As expected, they show the same trend with PPIs (Table S2). The correlation coefficient between degrees and ProTINA scores is 0.208 for ‘Rapamycin’, while that for betweenness values is 0.079 (Figure 5a, b).

**Figure 5.**
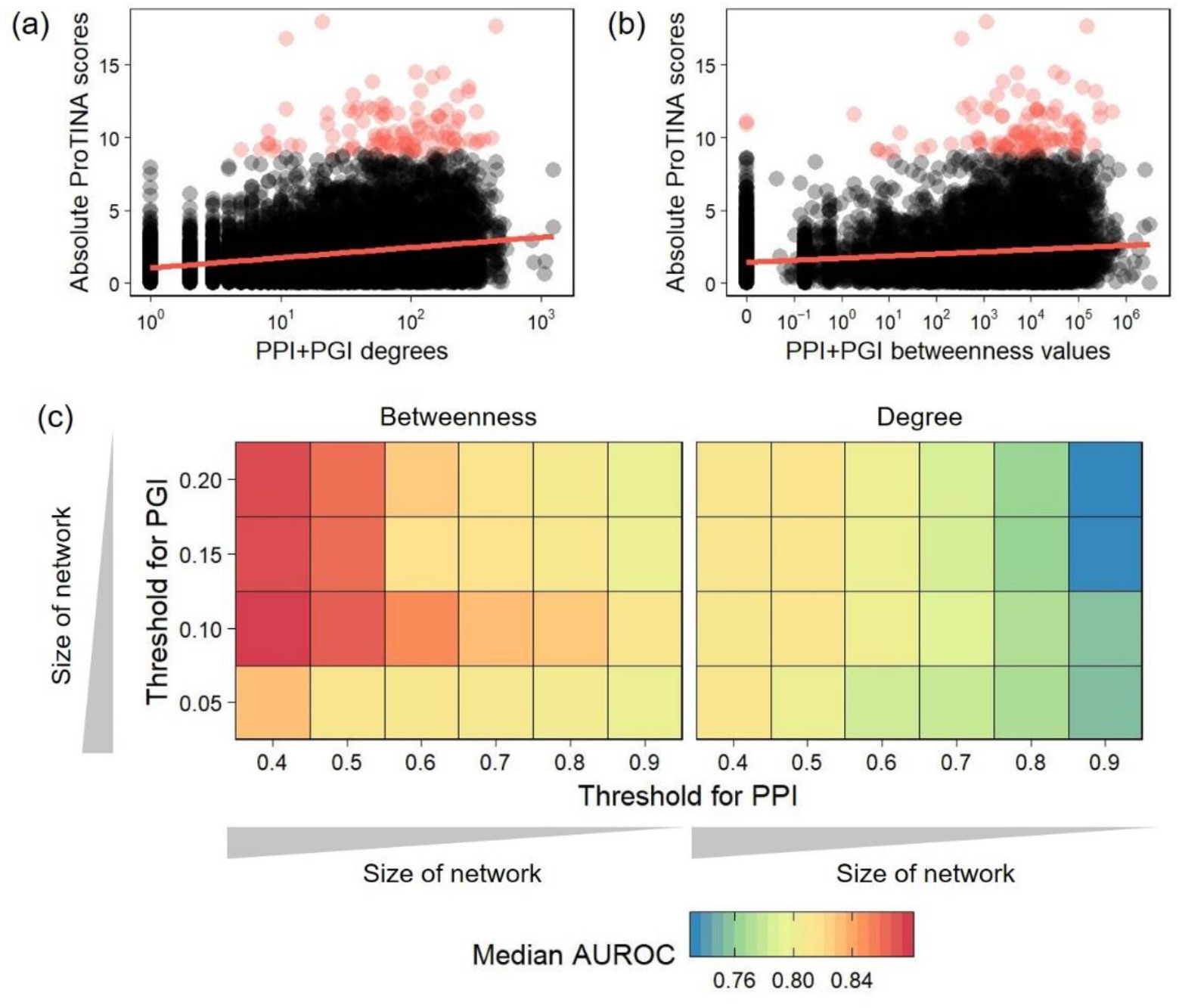
(a) PPI+PGI degree or (b) betweenness values were compared with associated ProTINA scores for ‘Rapamycin’. The correlation coefficient for absolute ProTINA scores versus degrees and absolute ProTINA scores versus betweenness values is 0.208 and 0.079, respectively. Red points refer to the top 100 proteins scored by ProTINA. (c) The degree and betweenness values were used as measures to predict drug targets, and the median AUROC values were calculated for each prediction. The axes refer to the confidence thresholds for PPI (x axis) and PGI (y axis) subnetworks.

Predicting drug targets by PPI+PGI degree or betweenness values results in higher median AUROC values than those for PPIs. In addition, for all thresholds of PPIs or PGIs applied to this analysis, betweenness values outperform degrees. The accuracy for PPI+PGI degrees ranges from 0.833 to 0.733 in terms of median AUROC values, and that for betweenness values ranges from 0.878 to 0.782 (Figure 5c). As the size of PPI or PGI subnetworks decreases, the median AUROC values for PPI+PGI degrees also decreases. PPI+PGI betweenness values have the same behavior as the size of PPIs changes, while for PGIs the trend is less evident. PGI10 has the best performance in parallel comparisons. In summary, betweenness values well predict drug targets and show even higher accuracy than ProTINA.

### Missing information in network topology can be covered by differential expression

Topological features, degree or betweenness values, have shown high prediction accuracy without taking gene expression into account. Our permutation tests have also indicated that network data has more effects in ProTINA’s performance than gene expression data. What’s more, ProTINA has much better performance than differential expression (DE) analysis on drug target prediction according to prior research [14]. All of the above has raised a question about whether gene expression data can help to predict drug targets. To address this concern, we compared DE analysis (adjusted *p*-values, Materials and Methods) with two other target prediction methods in terms of their performance on each drug: PPI+PGI betweenness and ProTINA.

We calculated AUROC values for all three methods using the same network setup, PPI09 and PGI20 (Figure 6). For most drugs, such as ‘Mitomycin C’ or ‘Cycloheximide’, PPI+PGI betweenness and ProTINA have close AUROC values, and they outperform DE analysis. Consistent behaviors between ProTINA and PPI+PGI further indicates the impact of network topology on ProTINA’s accuracy. In contrast to these drugs, DE has much higher prediction accuracy than PPI+PGI or ProTINA for ‘Monastrol’ (the AUROCs are 0.998, 0.771 and 0.555, respectively). This means that DE analysis of gene expression data can capture information missing in network topology, and that it is necessary to include gene expression data for drug target inference and improvement of accuracy.

**Figure 6.**
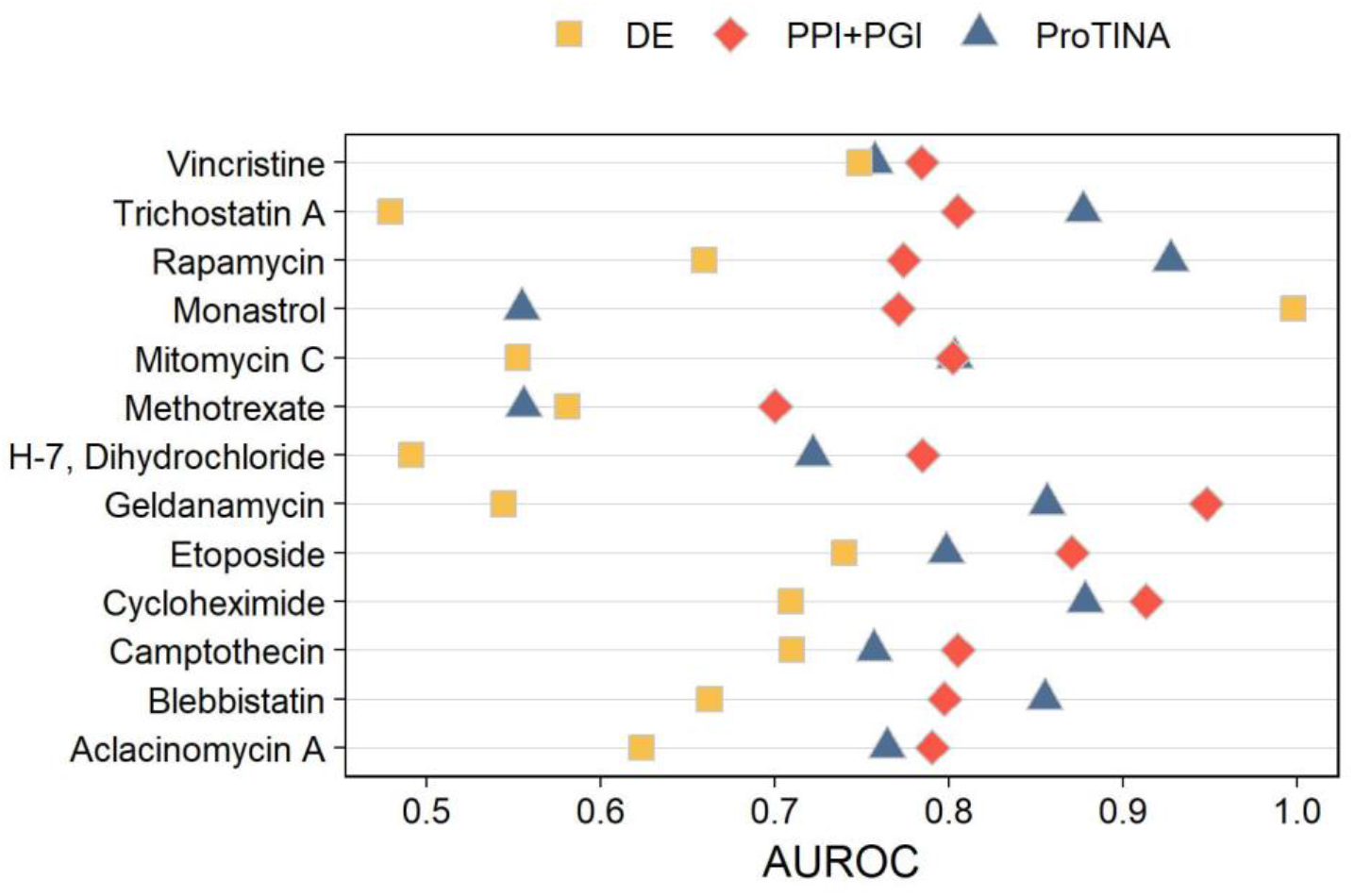
AUROC values of each drug obtained from three different methods: differential expression (DE) analysis by adjusted *p*-values, betweenness values from the combination of PPI09 and PGI20 (PPI+PGI) and ProTINA analysis by PPI09 and PGI20.

### A novel algorithm that combines network topology and DE analysis for target inference

To better combine network topology and DE analysis and improve inference accuracy, we suggest TREAP (<u>t</u>arget inference by <u>r</u>anking b<u>e</u>tweenness values and <u>a</u>djusted *<u>p</u>*-values) to predict drug targets. There are three steps for this algorithm. The first step is to calculate PPI+PGI betweenness values and obtain adjusted *p*-values from DE analysis. For gene expression profiles with multiple timepoints, DE analysis can be performed per time point or across all timepoints. The second step is to calculate the ranks of genes by sorting betweenness values and adjusted *p*-values, respectively. Genes with high betweenness values or low adjusted *p*-values are scored with high ranks. The third step is to generate final scores by summing up the ranks from both metrics for each gene. Genes with higher scores are more likely to be targets for associated drugs.

TREAP was tested by three different gene expression profiles: (i) DP14 [35], (ii) human HepG2 cells treated with genotoxic or non-genotoxic chemicals, referred to as HepG2 in this work [36] and (iii) mouse pancreatic cell lines treated with chromatin-targeting compounds, referred to as MP [37]. Human and mouse PPIs were obtained from STRING [28], and PGIs were obtained from Regulatory Circuits [40], TRRUST (version 2) [41] and RegNetwork [42]. AUROCs were calculated by comparing scored genes with known targets for each test as a measurement of accuracy.

TREAP shows stable performance and maintains high accuracy for all datasets tested in this study (median AUROCs > 0.800, Figure 7). While TREAP takes significantly less computation time than ProTINA, it has higher median AUROCs when compared with ProTINA under the same dataset (Figure 7). For DP14, the median AUROC of TREAP is 0.850, higher than that of ProTINA, 0.799 (*p*-value = 0.11). Notice that it is also higher than using PPI+PGI betweenness values alone, which is 0.798. TREAP significantly outperforms ProTINA in HepG2. The median AUROC is significantly improved from 0.739 to 0.801, with a *p*-value of 0.0002. For MP, TREAP and ProTINA have close median AUROCs as 0.806 and 0.799, respectively (*p*-value = 0.39). By integrating betweenness values and adjusted *p*-values to represent both network topology and DE analysis, TREAP is comparable with and sometimes better than ProTINA in accuracy for all datasets analyzed in this work. In addition, we performed 1000 permutation tests on TREAP by randomizing the adjusted *p*-values for each dataset. Different from ProTINA, TREAP is significant when compared with permutation tests on the adjusted *p*-values from DP14, with the one-tailed *p*-value as 0.007 (Figure S6). For HepG2, the one-tailed *p*-value is 0.058, while for MP, TREAP is less significant and shows a one-tailed *p*-value as 0.314.

**Figure 7.**
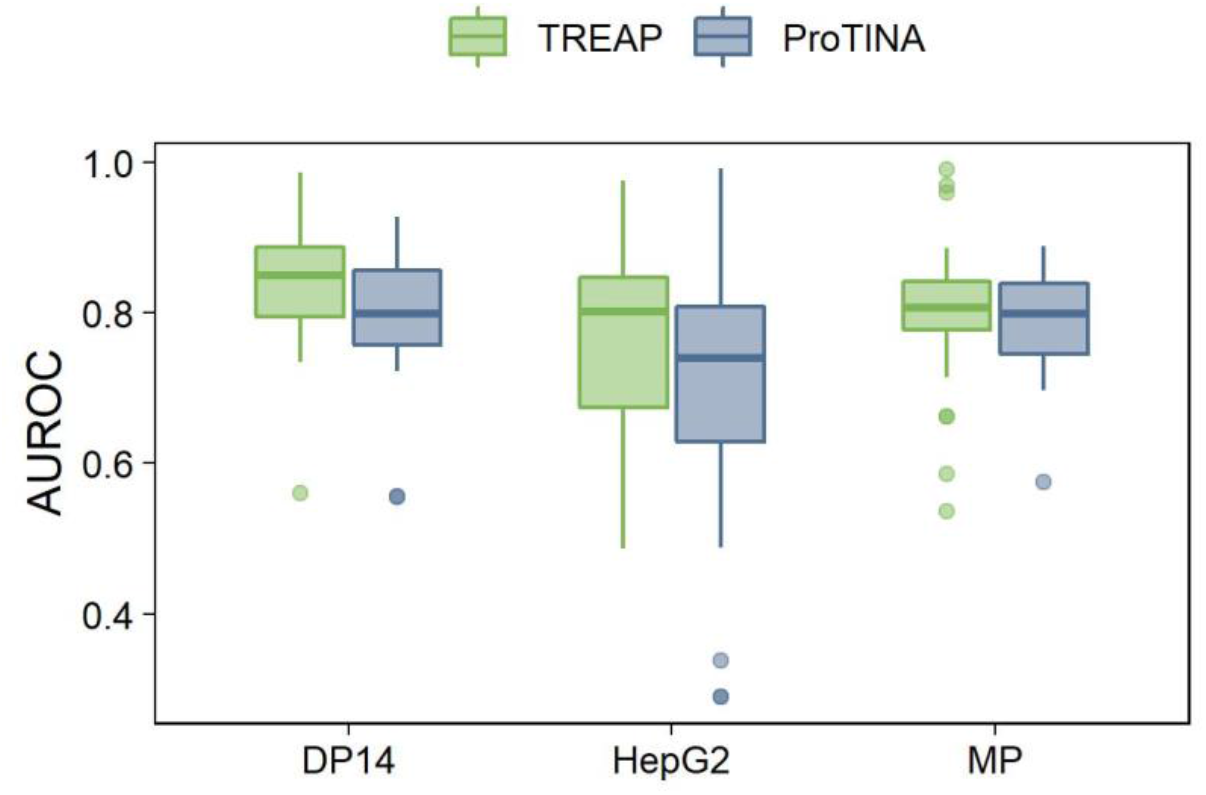
AUROC values of TREAP and ProTINA predictions for different gene expression profiles: human lymphoma cells (DP14), human liver cancer cells (HepG2) and mouse pancreatic cells (MP). (*p*-values = 0.11, 0.0002 and 0.39, respectively)

## Discussion

Our analyses have shown that, even though ProTINA requires both gene expression and network data for inputs, network data predominantly determines accuracy of drug target inference. What’s more, the cell/tissue type or size of a network has limited impact on ProTINA’s accuracy, while topology, especially the degree value, affects the performance of ProTINA more.

However, ProTINA has two limitations due to the reliance on network topology alone and the connection with protein degrees. First, PPIs, a major part of network data, have a known bias toward protein abundance [33, 47]. It has been reported that interactions obtained from high-throughput experiments have a correlation between the protein degree and abundance. Second, we have shown that differential analysis of gene expression data can uncover meaningful information missing in network topology.

To address these two limitations, we suggested a new algorithm, TREAP, which combines protein betweenness values and adjusted *p*-values, representing information from both sources of network topology and DE analysis, to predict drug targets. We chose betweenness values because they are less sensitive to network sizes and more accurate than degrees in target prediction. TREAP shows more consistent performance than ProTINA when tested by different gene expression profiles and maintains a median AUROC above 0.800. In addition, TREAP takes significantly less computation time than ProTINA, and its simplicity makes it more tractable to users who are not experts in systems and network biology. It is also flexible in dealing with samples of limited or multiple timepoints as adjusted *p*-values can be calculated per timepoint or across all timepoints based on user’s needs. However, ProTINA needs at least two timepoints to fully take advantage of the algorithm [14]. Currently, betweenness values and adjusted *p*-values are weighted equally for TREAP. Future work should focus on better balancing both types of data and trying other scoring methods to improve prediction accuracy.

TREAP is presented here as an alternative approach to ProTINA, but it is worth emphasizing advantages specific to each algorithm. As stated above, TREAP is significantly faster, and the algorithm is not complex, enabling users from several branches of research to access the tool and understand the findings. ProTINA, however, is a mechanistically derived algorithm, which allows users with expertise in computational biology to dive deeper into the possible mechanisms of a drug’s activity. The accuracy of their predictions is similar when measured using the AUROC, but the permutation tests presented here suggest that TREAP is more likely to use drug-specific gene expression to make a more accurate prediction.

TREAP and its derivatives have potential in a variety of applications for drug innovation. First, it can assist in selection of drug candidates and serve as a preliminary test of the efficacy or safety by connecting with databases for functional annotations, e.g. Gene Ontology [48, 49]. Studying predicted targets can help exclude poorly targeting drug candidates or those causing severe damage to biological systems. Second, the algorithm can be applied to drug repurposing by exploring published datasets characterizing drug treatments, assuming that a pair of drugs sharing the same group of predicted targets can be used to treat the same disease. Last but not the least, the algorithm can help to discover disease mechanisms [14]. Similar to drug treatments, diseases can also be treated as a type of perturbation to the biological system of interest. Predicting disease targets may assist in identifying key components of disease mechanisms and pathology, which is crucial for innovations in disease treatment [10].

## Author Contributions

conceptualization, J.E.S. and M.W.; methodology, M.W.; software, M.W.; formal analysis, M.W.; investigation, M.W. and E.M.; writing–original draft preparation, M.W.; writing–review and editing, J.E.S., E.M. and N.H.; visualization, M.W.; supervision, J.E.S.

## Supplementary Materials

Figure S1: Comparing the accuracy of ProTINA and DeMAND; Figure S2: The number of interactions under different thresholds for PPIs and PGIs; Figure S3: Performance of ProTINA on DP14 using PPIs and PGIs of different sizes; Figure S4: Performance of ProTINA on DP14 using PGIs from different cell/tissue types; Figure S5: Predicting drug targets by degree or betweenness values of PGIs; Figure S6: Median AUROCs of permutation tests on TREAP. Table S1: Reference drug targets for each analysis; Table S2: Correlation coefficients between degree or betweenness values and ProTINA scores for each drug; Table S3: AUROCs of each drug for ProTINA, topological features and TREAP; Table S4: Degree and betweenness values of proteins under different thresholds.

## References

1. Schuhmacher, A., O. Gassmann, and M. Hinder, Changing R&D models in research-based pharmaceutical companies. J Transl Med, 2016. 14(1): p. 105.

2. Scannell, J.W., et al., Diagnosing the decline in pharmaceutical R&D efficiency. Nat Rev Drug Discov, 2012. 11(3): p. 191–200.

3. Pushpakom, S., et al., Drug repurposing: progress, challenges and recommendations. Nat Rev Drug Discov, 2019. 18(1): p. 41–58.

4. Paul, S.M., et al., How to improve R&D productivity: the pharmaceutical industry’s grand challenge. Nat Rev Drug Discov, 2010. 9(3): p. 203–14.

5. DiMasi, J.A., H.G. Grabowski, and R.W. Hansen, Innovation in the pharmaceutical industry: New estimates of R&D costs. J Health Econ, 2016. 47: p. 20–33.

6. Cook, D., et al., Lessons learned from the fate of AstraZeneca’s drug pipeline: a five-dimensional framework. Nat Rev Drug Discov, 2014. 13(6): p. 419–31.

7. Hay, M., et al., Clinical development success rates for investigational drugs. Nat Biotechnol, 2014. 32(1): p. 40–51.

8. Hwang, T.J., et al., Failure of Investigational Drugs in Late-Stage Clinical Development and Publication of Trial Results. JAMA Intern Med, 2016. 176(12): p. 1826–1833.

9. Harrison, R.K., Phase II and phase III failures: 2013-2015. Nat Rev Drug Discov, 2016. 15(12): p. 817–818.

10. Spagnolo, P. and T.M. Maher, Clinical trial research in focus: why do so many clinical trials fail in IPF? Lancet Respir Med, 2017. 5(5): p. 372–374.

11. Naci, H. and J.P. Ioannidis, How good is “evidence” from clinical studies of drug effects and why might such evidence fail in the prediction of the clinical utility of drugs? Annu Rev Pharmacol Toxicol, 2015. 55: p. 169–89.

12. Gashaw, I., et al., What makes a good drug target? Drug Discovery Today, 2012. 17: p. S24–S30.

13. Chua, H.N. and F.P. Roth, Discovering the targets of drugs via computational systems biology. J Biol Chem, 2011. 286(27): p. 23653–8.

14. Noh, H., J.E. Shoemaker, and R. Gunawan, Network perturbation analysis of gene transcriptional profiles reveals protein targets and mechanism of action of drugs and influenza A viral infection. Nucleic Acids Res, 2018.

15. Lamb, J., et al., The Connectivity Map: using gene-expression signatures to connect small molecules, genes, and disease. Science, 2006. 313(5795): p. 1929–35.

16. Ganter, B., et al., Development of a large-scale chemogenomics database to improve drug candidate selection and to understand mechanisms of chemical toxicity and action. J Biotechnol, 2005. 119(3): p. 219–44.

17. Wang, M., C. Tang, and J. Chen, Drug-Target Interaction Prediction via Dual Laplacian Graph Regularized Matrix Completion. Biomed Res Int, 2018. 2018: p. 1425608.

18. Vertes, A., et al. Inferring Mechanism of Action of an Unknown Compound from Time Series Omics Data. 2018. Cham: Springer International Publishing.

19. Wolpaw, A.J., et al., Modulatory profiling identifies mechanisms of small molecule-induced cell death. Proc Natl Acad Sci U S A, 2011. 108(39): p. E771–80.

20. Iorio, F., et al., Transcriptional data: a new gateway to drug repositioning? Drug Discov Today, 2013. 18(7-8): p. 350–7.

21. Rush, S.T.A. and D. Repsilber, Capturing context-specific regulation in molecular interaction networks. BMC Bioinformatics, 2018. 19(1): p. 539.

22. Woo, J.H., et al., Elucidating Compound Mechanism of Action by Network Perturbation Analysis. Cell, 2015. 162(2): p. 441–451.

23. Chen, E.Y., et al., Expression2Kinases: mRNA profiling linked to multiple upstream regulatory layers. Bioinformatics, 2012. 28(1): p. 105–11.

24. Koido, M., et al., InDePTH: detection of hub genes for developing gene expression networks under anticancer drug treatment. Oncotarget, 2018. 9(49): p. 29097–29111.

25. Ji, X., J.M. Freudenberg, and P. Agarwal, Integrating Biological Networks for Drug Target Prediction and Prioritization. Methods Mol Biol, 2019. 1903: p. 203–218.

26. Cosgrove, E.J., et al., Predicting gene targets of perturbations via network-based filtering of mRNA expression compendia. Bioinformatics, 2008. 24(21): p. 2482–90.

27. Failli, M., J. Paananen, and V. Fortino, Prioritizing target-disease associations with novel safety and efficacy scoring methods. Sci Rep, 2019. 9(1): p. 9852.

28. Szklarczyk, D., et al., STRING v11: protein-protein association networks with increased coverage, supporting functional discovery in genome-wide experimental datasets. Nucleic Acids Res, 2019. 47(D1): p. D607–D613.

29. Cahan, P., et al., CellNet: network biology applied to stem cell engineering. Cell, 2014. 158(4): p. 903–915.

30. Santolini, M. and A.L. Barabasi, Predicting perturbation patterns from the topology of biological networks. Proc Natl Acad Sci U S A, 2018. 115(27): p. E6375–E6383.

31. Zhu, M., et al., The analysis of the drug–targets based on the topological properties in the human protein–protein interaction network. Journal of Drug Targeting, 2009. 17(7): p. 524–532.

32. Lopes, T.J., et al., Identifying problematic drugs based on the characteristics of their targets. Front Pharmacol, 2015. 6: p. 186.

33. Ackerman, E.E., et al., Network-Guided Discovery of Influenza Virus Replication Host Factors. MBio, 2018. 9(6).

34. Feng, Y., Q. Wang, and T. Wang, Drug Target Protein-Protein Interaction Networks: A Systematic Perspective. Biomed Res Int, 2017. 2017: p. 1289259.

35. Bansal, M., et al., A community computational challenge to predict the activity of pairs of compounds. Nat Biotechnol, 2014. 32(12): p. 1213–22.

36. Magkoufopoulou, C., et al., A transcriptomics-based in vitro assay for predicting chemical genotoxicity in vivo. Carcinogenesis, 2012. 33(7): p. 1421–9.

37. Kubicek, S., et al., Chromatin-targeting small molecules cause class-specific transcriptional changes in pancreatic endocrine cells. Proc Natl Acad Sci U S A, 2012. 109(14): p. 5364–9.

38. Gautier, L., et al., affy—analysis of Affymetrix GeneChip data at the probe level. Bioinformatics, 2004. 20(3): p. 307–315.

39. Ritchie, M.E., et al., limma powers differential expression analyses for RNA-sequencing and microarray studies. Nucleic Acids Research, 2015. 43(7): p. e47–e47.

40. Marbach, D., et al., Tissue-specific regulatory circuits reveal variable modular perturbations across complex diseases. Nat Methods, 2016. 13(4): p. 366–70.

41. Han, H., et al., TRRUST v2: an expanded reference database of human and mouse transcriptional regulatory interactions. Nucleic Acids Res, 2018. 46(D1): p. D380–D386.

42. Liu, Z.P., et al., RegNetwork: an integrated database of transcriptional and post-transcriptional regulatory networks in human and mouse. Database (Oxford), 2015. 2015.

43. Csardi, G. and T. Nepusz, The Igraph Software Package for Complex Network Research. InterJournal, 2005. Complex Systems: p. 1695.

44. Kuhn, M., et al., STITCH: interaction networks of chemicals and proteins. Nucleic acids research, 2008. 36(Database issue): p. D684–D688.

45. Szklarczyk, D., et al., STITCH 5: augmenting protein–chemical interaction networks with tissue and affinity data. Nucleic acids research, 2016. 44(D1): p. D380–D384.

46. Robin, X., et al., pROC: an open-source package for R and S+ to analyze and compare ROC curves. BMC Bioinformatics, 2011. 12(1): p. 77.

47. Ivanic, J., et al., Influence of protein abundance on high-throughput protein-protein interaction detection. PLoS One, 2009. 4(6): p. e5815.

48. The Gene Ontology, C., The Gene Ontology Resource: 20 years and still GOing strong. Nucleic Acids Res, 2019. 47(D1): p. D330–D338.

49. Ashburner, M., et al., Gene ontology: tool for the unification of biology. The Gene Ontology Consortium. Nat Genet, 2000. 25(1): p. 25–9.

